# β-Glucan Reprograms Neutrophils to Induce Disease Tolerance Against Influenza A Virus

**DOI:** 10.1101/2024.09.02.610822

**Authors:** Nargis Khan, Raphael Chevre, Sarah Sun, Mina Sadeghi, Erwan Pernet, Andrea Herrero, Alexander Grant, Jeffrey Downey, Luis B. Barreiro, Bryan G Yipp, Oliver Soehnlein, Maziar Divangahi

**Author notes:** Lead Contact: Maziar Divangahi.

## Abstract

Disease tolerance is an evolutionarily conserved host defence strategy that preserves tissue integrity and physiology without affecting pathogen load. Unlike host resistance, the mechanisms underlying disease tolerance remain poorly understood. In the present study, we investigated whether an adjuvant (β-glucan) can reprogram innate immunity to provide protection against Influenza A virus (IAV) infection. Here we observe that β-glucan treatment reduced the morbidity and mortality against IAV infection, independent of host resistance (viral load). Increased survival of β-glucan treated mice against IAV is associated with the accumulation of neutrophils via RoRγt^+^ T cells in the lung tissue. Using gain-and-loss-of-function approaches, we demonstrate that β- glucan reprogrammed neutrophils are essential for promoting disease tolerance, limiting pulmonary tissue damage, and enhancing survival against IAV infection. β-glucan treatment promotes granulopoiesis in a type 1 interferon-dependent manner that leads to the generation of a unique subset of neutrophils, which are less mature with higher mitochondrial mass utilizing mitochondrial oxidative (OXPHOS) metabolism. Collectively, our data indicate that β-glucan reprograms hematopoietic stem cells (HSCs) to generate neutrophils with a novel “regulatory” function, which is required for promoting disease tolerance and maintaining lung tissue integrity against viral infection.

## INTRODUCTION

It is increasingly understood that host defence strategies against infectious diseases are comprised of both *host resistance* and *disease tolerance*. Host resistance is the ability to prevent invasion or to eliminate the pathogen, while disease tolerance is defined by limiting the tissue damage caused by the pathogen and/or the immune response^1, 2, 3^. Unlike resistance, disease tolerance does not necessarily exert direct effects on pathogen growth or survival. Disease tolerance plays a critical role in pulmonary infections *via* promoting tissue repair mechanisms for maintaining lung tissue integrity and functions. For instance, we have recently shown how pulmonary macrophages^4^ or NK cells^5^ can promote disease tolerance against influenza A virus (IAV) infection and enhance host survival independent of lung viral replication. However, the mechanism of how we can harness the power of disease tolerance against infectious diseases remains largely unknown.

The capacity of innate immune cells to maintain memory (termed trained immunity) has revealed an important and unrecognized property of innate immune responses that can be targeted to enhance host defense against infectious diseases. While there has been mounting evidence showing that the innate immune system can confer long-term functional reprogramming in either homologous or heterologous infections by mainly boosting host resistance^6, 7^, little is known about the contribution of innate immune memory responses in promoting disease tolerance against infectious diseases. Considering the relatively short lifespan of innate cells, we and others have demonstrated that the maintenance of innate immune memory cells relies on reprogramming the bone marrow hematopoietic stem cells (HSCs) that give rise to progenitor and mature innate immune cells^8^. For instance, β-glucan (a fungal cell wall component) reprograms HSCs and generates trained monocytes promoting host resistance against chronic *Mycobacteria tuberculosis* infection, which remarkably enhances host survival^9^. Similarly, β-glucan has been shown to protect against various acute bacterial infections, fungi, or tumours^10, 11^, but its role against pulmonary viral infection is unclear.

The severity of pulmonary viral infections is often a reflection of lung damage, which is the dominant feature of fatal outcomes^12^. Immuno-pathology is attributed to an over-exuberant inflammatory response in the airways and lungs, frequently initiated by neutrophil recruitment. Neutrophils are the most abundant cell types, representing 50 to 70% of the total circulatory leukocytes with a short lifespan (∼24h)^13^. At steady-state, neutrophils contribute to tissue repair and homeostasis by phagocytosing necrotic cells and producing resolvins and protectins^14^. It is well established that neutrophils’ antimicrobial functions are essential against infections as neutrophil disorders are associated with recurrent bacterial or fungal infections^15^. However, recent studies have shown that the functional spectrum of neutrophils is extremely diverse, ranging from the ability to program alveolar macrophages in the developing lung^16^ to limiting tissue damage and promoting wound healing following injury or infection^17, 18, 19^ . Thus, dissecting the cellular and molecular mechanisms imprinting neutrophil heterogeneity may provide an opportunity for targeting them during infections.

Considering β-glucan provides protection against bacterial or fungal infections, we sought to investigate the impact of β-glucan during IAV infection. Here we show that β-glucan reprogrammes HSCs and promotes granulopoiesis via type I IFN signaling, which gives rise to a unique subset of regulatory neutrophils. These neutrophils exhibit a less mature phenotype with an altered metabolic program. Unlike their classical role in host resistance, β-glucan-mediated regulatory neutrophils promote disease tolerance against IAV via limiting the lung pathology independent of viral replication.

## RESULTS

### β-glucan promotes disease tolerance against Influenza A virus infection

β-glucan has been known as an inducer of trained immunity^20^. Recently, we demonstrated that β- glucan trains monocytes and provides protection against TB^9^. However, the potential contribution of β-glucan in host defense against pulmonary viral infection remains largely unknown. To investigate this, we treated the C57BL/6 mice with β-glucan intraperitoneally (i.p.), and after 7 days, mice were infected with influenza A virus (IAV) (Fig 1A). Mice treated with β-glucan showed significantly reduced morbidity and increased survival against IAV (Fig 1B-C). Increased survival of β-glucan-treated mice infected with IAV was independent of the host resistance, as there was no difference in viral load on days 1, 3, and 6 post-IAV infection (Fig. 1D). However, β-glucan-treated mice infected with IAV showed remarkably reduced immunopathology compared to control infected mice (Fig. 1E). To assess the longevity of β-glucan-mediated protection against IAV, mice were treated with β-glucan 30 days prior to IAV infection (Fig 1F).

**Figure 1.**
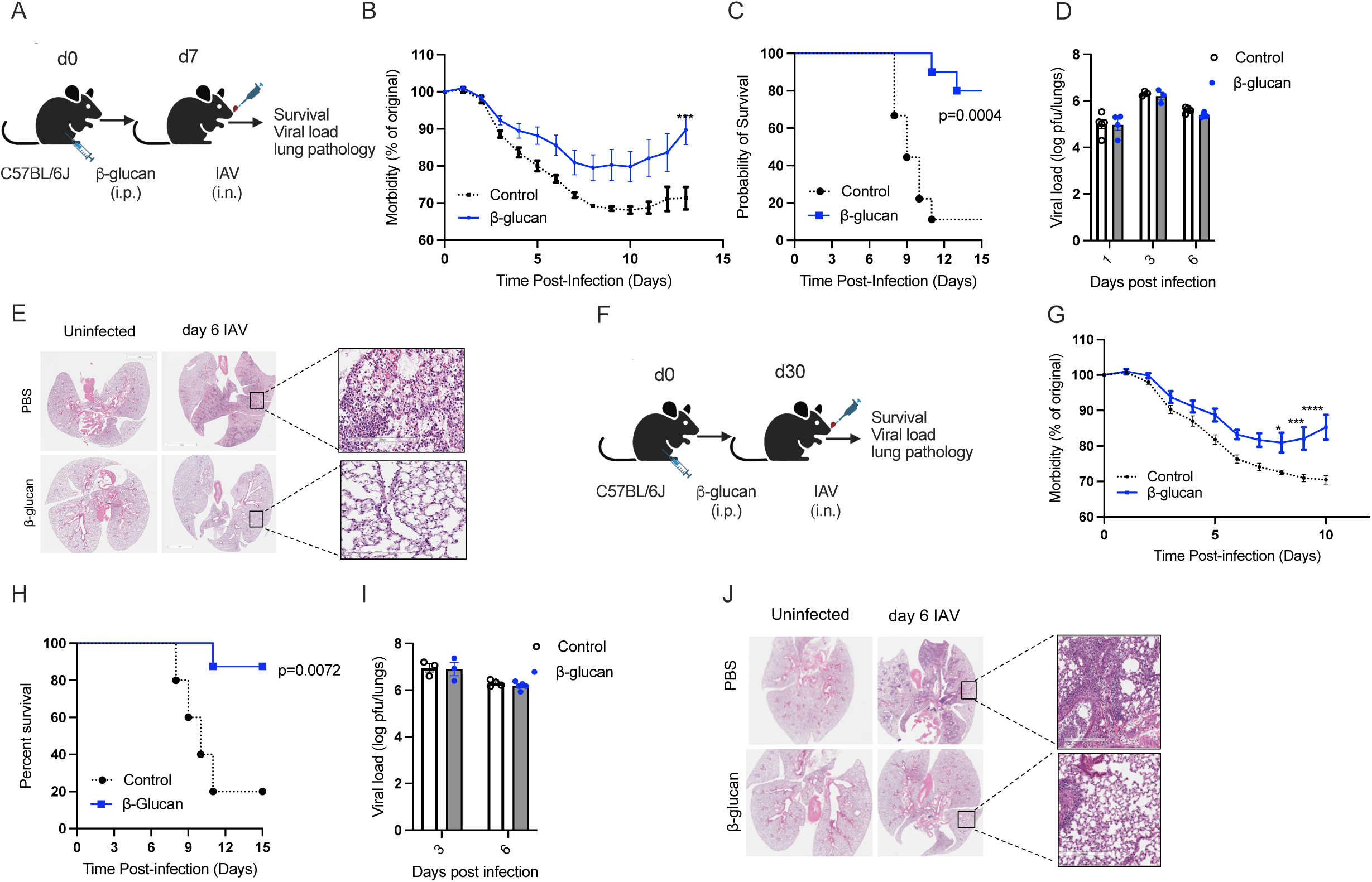
β-glucan treatment induces disease tolerance against IAV. (A-C)) Mice were infected with IAV (lethal dose) at day 7 post -β-glucan (i.p.) treatment (A). Morbidity (B) and survival (C) were monitored over time. (D-E) Mice were infected with a sublethal dose at day seven post-glucan treatment. Viral load was measured at various time points post-infection (D). Representative micrographs of lungs from β-glucan treated mice stained with hematoxylin and eosin day 6 post-IAV infection (Scale bar represents 200 uM (Magnified image) (E). (F-H) Mice were treated with β-glucan (i.p.) followed by IAV infection with lethal (survival and morbidity) or sublethal dose (viral load and pathology) at day 30 post -β-glucan (i.p.) treatment. Morbidity (G) and Survival (H) were monitored over time. Viral load at days 3 and 6 post-infection (I). Representative micrographs of lungs from β-glucan treated mice stained with hematoxylin and eosin at day six post-IAV infection. (Scale bar represents 200 uM (Magnified image) (J). Data were analyzed using one-way ANOVA followed by Tukey’s multiple comparisons test or two- way ANOVA followed by Sidak’s multiple comparisons tests. Survival was monitored by a log- rank test. * *p*<.05, *** *p*<0.001, **** *p*<0.0001.

Similar to the β-glucan short-treatment (7 days), the protective impact of the β-glucan long-treatment on morbidity and mortality was maintained following IAV infection (Fig. 1G-I). This enhanced host defense was independent of host resistance but with reduced lung immunopathology (Fig 1 J). Collectively, these data indicate that β-glucan treatment protects IAV by promoting disease tolerance rather than enhancing anti-viral response.

### β-glucan treatment increases the recruitment of neutrophils into the lung tissue

The ability of β-glucan to enhance host immune response is well documented^9, 21^. Recently, we demonstrated that β-glucan enhances host resistance against *Mtb*, and the protection was mediated by trained monocytes and macrophages^9^. In contrast to *Mt*b, the protection of β-glucan against IAV was independent of pulmonary viral load, but reduced immunopathology suggests β-glucan may regulate disease tolerance against IAV (Fig. 1A-J). Interestingly, immunophenotyping of β- glucan-treated mice showed a significant increase in the frequency and absolute number of neutrophils in the lungs, which peaked on day four and returned to the basal level by day seven (Fig. 2A-D; SFig. 1A-C Gating Strategy). However, the differences in the other pulmonary myeloid or lymphoid cells, such as monocytes/macrophages or T cells, were minimal (S. Fig. 2 A, B). Furthermore, this neutrophilia was not limited to the lungs of β-glucan-treated mice, as the number of neutrophils was also increased in the blood, spleen, and peritoneal cavity (S Fig. 2C- G).

**Figure 2.**
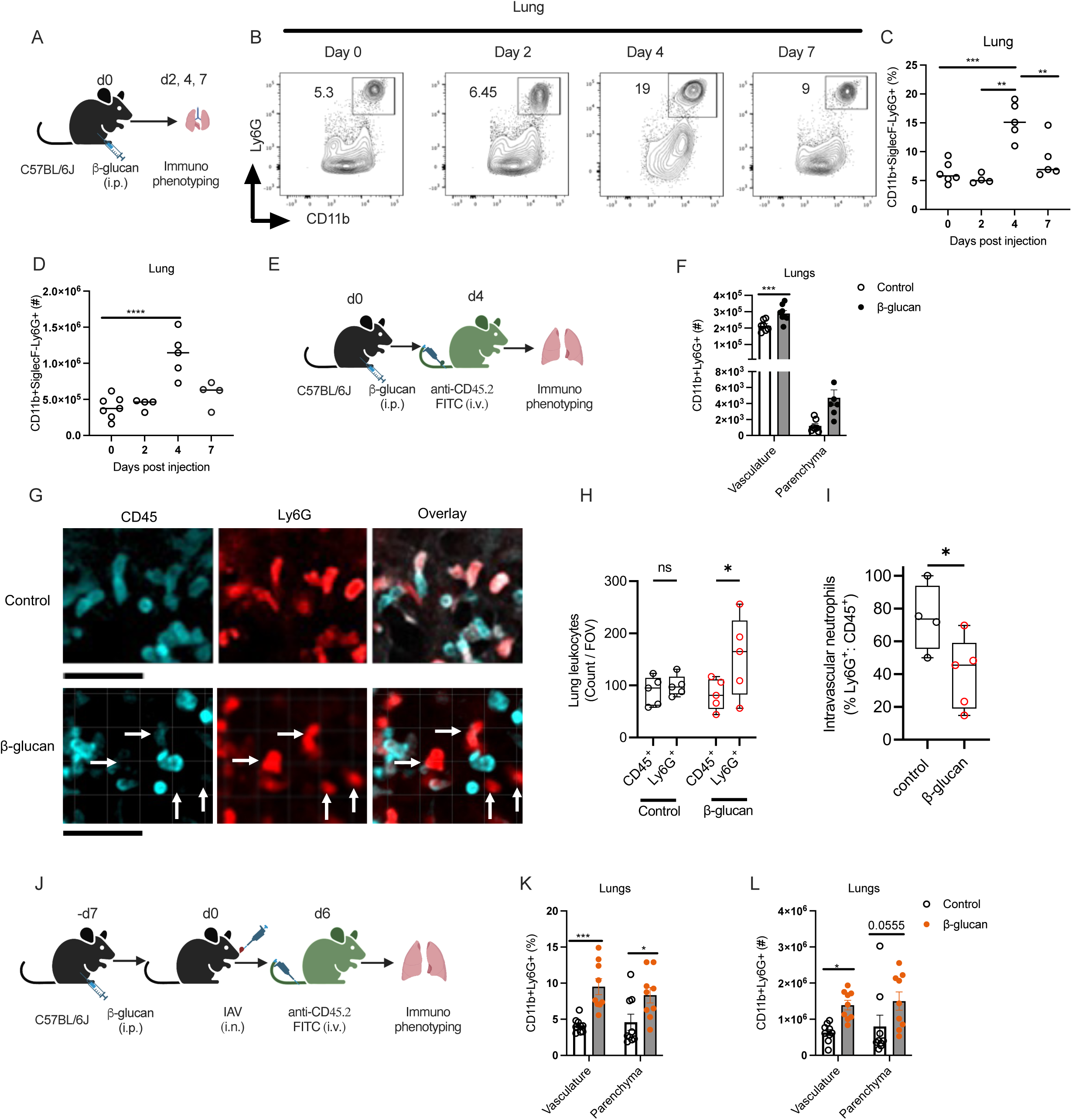
β-glucan treatment increases the recruitment of neutrophils to the lungs. (A-D) Mice were treated with β-glucan. Immune cells were assessed in the BM. Representative FACS plots (B), frequency (C), and total cell counts (D) of neutrophils in the lungs at days 2, 4, and 7 posts-β-glucan treatment. (E-F) Intravascular staining at day four post-β-glucan treatment. Total cells count of neutrophils in the vasculature and parenchyma of lungs (F). (G) Representative lung confocal intravital microscopy images comparing Ly6G-TdTom mice receiving saline i.p. versus mice treated with β-glucan i.p. Intravenous fluorescently conjugated anti-CD45 monoclonal antibody was used to mark intravascular leukocytes. Arrows highlight examples of Ly6G+CD45- cells. (H) Visualized cells from the lung intravital were quantified by expression of either CD45 or Ly6G. All scale bars = 50µm. (I), The percentage of intravascular neutrophils in lung imaging was quantified as Ly6G+CD45+ /total Ly6G+. (J) Intra-vasculature staining at day 6 post- IAV infection. Frequency (K) and total cell counts (L) of neutrophils in the vasculature and parenchyma of lungs day 6 post-IAV infection. Data were analyzed using one-way ANOVA followed by Tukey’s multiple comparisons test or two-way ANOVA followed by Sidak’s multiple comparisons tests. * p<0.05, ** p<0.01, *** p<0.001, **** p<0.0001.

Recent studies in neutrophil dynamics suggest that the trafficking of neutrophils from the vasculature into tissue is critical in defining their functions^22^. The intravascular staining showed that number of neutrophils was significantly increased in both the lung vasculature as well as the parenchyma of mice treated with β-glucan (Fig 2E, F; SFig. 2H). To directly investigate the impact of β-glucan on lung neutrophil recruitment and localizing *in vivo*, pulmonary confocal intravital was performed in Ly6G-TdTom mice. Intravenously administered fluorescently conjugated anti- CD45 monoclonal antibodies were used to differentiate vascular neutrophils from non-vascular neutrophils. Corroborating the flow cytometry data, we found that β-glucan induced neutrophil recruitment, but these neutrophils were significantly less vascular (Fig. 2 H, I). Importantly, the intravascular staining also showed that both the frequency and absolute numbers of neutrophils in both the lung vasculature and parenchyma were also increased in β-glucan-treated mice after 6 days of infection with IAV (Fig. 2J-L). This was specific to neutrophils, as there was no difference in the frequency or absolute number of monocytes in the lungs (SFig.2I-L). Collectively, these results indicate that β-glucan increases the number of neutrophils in blood circulation and promotes the recruitment of neutrophils into the lung tissue.

### β-glucan promotes granulopoiesis in the bone marrow via the type I interferon signalling pathway

Recently, we have demonstrated that two doses of β-glucan expand hematopoietic stem cells (HSC) and promote myelopoiesis^9^. However, the increased number of neutrophils in the circulation of one dose of β-glucan treatment (Fig 2) suggests that the β-glucan reprogramming of HSC is biased towards granulopoiesis. Thus, we next investigated the effect of one dose of β- glucan on HSC expansion and the downstream progenitors (SFig. 1B, C; gating strategy). Both the frequency and absolute number of LKS (Lin-cKit+Sca-1+), MPP (LKS+CD150-CD48+), and GMP (Lin-cKit+CD34+CD16/32+) were significantly increased after β-glucan treatment (Fig. 3 A-D, SFig. 3 A-C). GMP is upstream of common monocyte progenitors (cMoPs) or granulocyte progenitors (GP) for producing neutrophils. Interestingly, we found increased GP (Lin-Sca-1- cKit+CD16/32+CD34+CD115-Ly6c+) in the BM with no differences in cMoPs (Lin-Sca-1- cKit+CD16/32+CD34+CD115+Ly6c-) suggesting that one-dose β-glucan promotes granulopoiesis (Fig 3 E: SFig. 3D-F) rather myelopoiesis following two-doses of β-glucan treatment. Increased GP in the BM correlated with the increased frequency and absolute number of neutrophils in the bone (Fig 3F-H).

**Figure 3.**
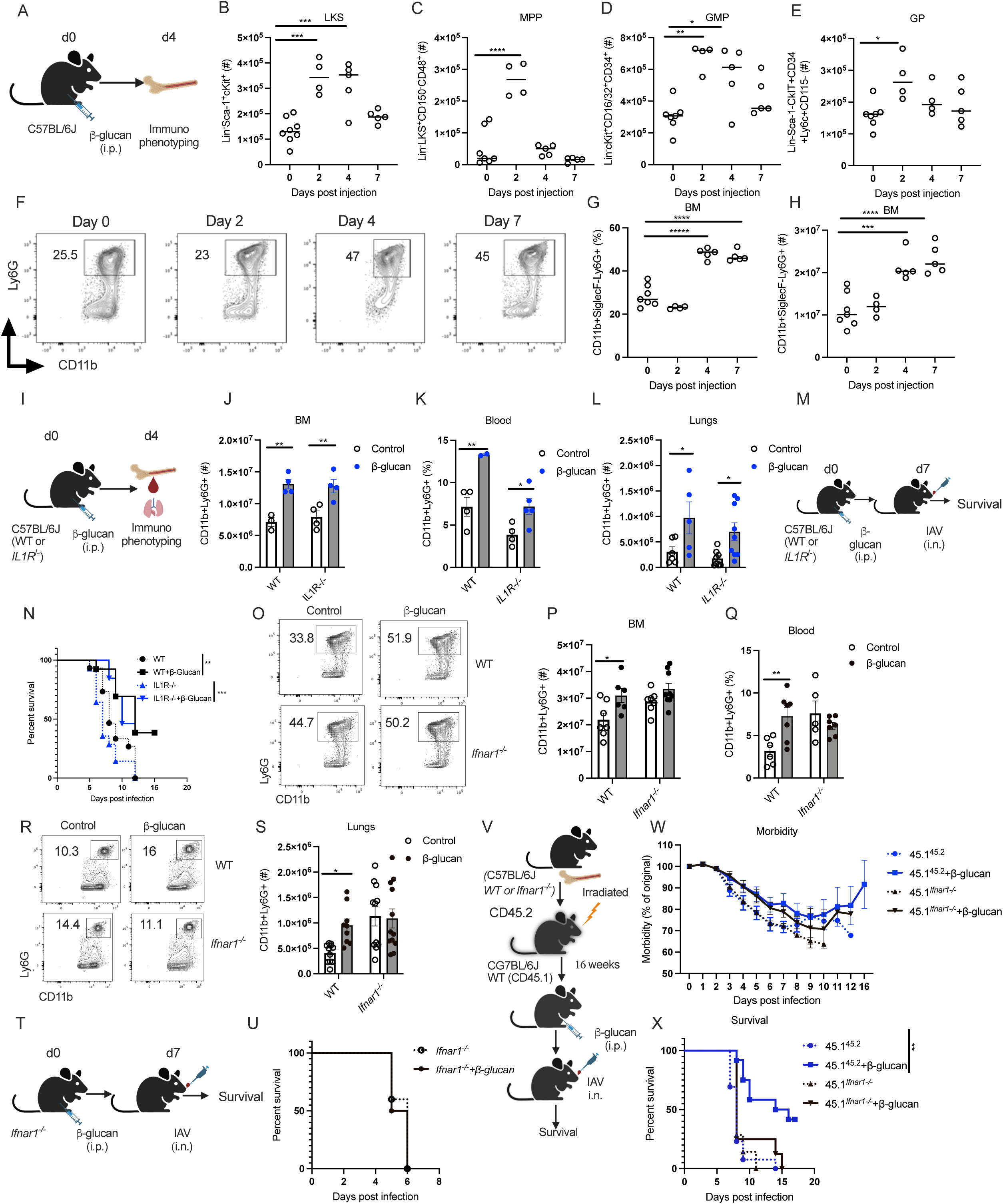
β-glucan increased granulopoiesis required type I interferon signalling. (A-C) Mice were treated with β-glucan. HSCs/progenitors and immune cells were assessed in the BM at days 2, 4, and 7 post-glucan treatment. Expansion of LKS (B), MPP (C), GMP (D) and GP (E) in the BM. (F-H) Representative FACS plots (F), frequency (G) and total cell counts of neutrophils (H) in the BM. (I-S) C57BL/6 (WT and *IL1R^-/-^* or *Ifnar1^-/-^*) mice were treated with β-glucan. Neutrophils in the BM (J), blood (K), and lungs (L) at day 4 post-glucan treatment. (M, N) Survival *of IL1R^-/-^* or WT mice after β-glucan treatment following IAV infection (lethal dose) at day 7. (O- R) WT and *Ifnar1^-/-^* mice were treated with β-glucan. Representative FACS plots (O), total cell counts (P) of neutrophils in the BM; frequency of neutrophils in the blood (Q); representative FACS plots (R); total cell counts of neutrophils (S) in the lungs at day 4 post β-glucan treatment. The number in the FACS plot indicates the viable frequency. (T, U) Survival of C57BL/6 (WT or *Ifnar1^-/-^*) mice after β-glucan treatment following IAV infection (lethal dose) at day 7. (V-X) Mouse chimera model (V). Morbidity (W) and survival (X) of CD45.1 chimeric mice reconstituted with *Ifnar1^-/-^*(CD45.2) BM after β-glucan treatment following IAV infection (lethal dose) on day 7. Data were analyzed using one-way ANOVA followed by Tukey’s multiple comparisons test or two-way ANOVA followed by Sidak’s multiple comparisons tests. Survival was monitored by a log-rank test. * p<0.05, ** p<0.01, *** p<0.001, **** p<0.0001.

Next, we investigated the signalling pathway involved in the β-glucan reprogramming of HSC towards granulopoiesis. Two doses of β-glucan reprogramming of HSC/myelopoiesis depend on IL1 signalling^9^. Thus, we next investigated whether IL1R signalling is involved in regulating granulopoiesis after one-dose β-glucan. After one dose of β-glucan-treatment, the number of neutrophils in the BM, blood and lungs was increased in IL1R^-/-^ mice (Fig. 3I-L). While there was no survival difference between IL1R^-/-^ and WT mice after a lethal dose of IAV infection, the survival of β-glucan-treated IL1R^-/-^ mice was also comparable to β-glucan-treated WT mice post-IAV infection (Fig. 3M, N). Unexpectedly, these data suggest that IL1 signalling is not required for the one-dose β-glucan-mediated granulopoiesis or protection against IAV.

Type I Interferon has been shown to promote anti-tumour solid immunity via switching the function of the tumour-associated neutrophils^23^. Therefore, we next investigated the potential role of type I IFN signalling in β-glucan training of neutrophils. Immuno-phenotyping of β-glucan treated *Ifnar1^-/-^* mice showed expansion of LKS, MPPs, and GMPs but failed to show an increase in GPs (SFig. 3I-P). Consequently, *Ifnar1^-/-^* mice showed no increased neutrophils in the BM, blood and lungs in response to β-glucan (Fig. 3O-S; SFig. 3Q, R) These data collectively indicate that β-glucan promotes the expansion of HSCs and GMPs independent of type I IFN signalling, but type I IFN signalling is required for driving the fate of GMPs towards GPs. Next, we infected *Ifnar1^-/-^* mice with IAV post 7 days of β-glucan treatment. We found that the *Ifnar1^-/-^* mice are equally susceptible and succumbed to death after IAV infection and this susceptibility was irrespective of β-glucan treatment (Fig. 3T, U). To determine the critical role of type I IFN signalling in the BM against IAV, we next generated chimeric mice by reconstituting the congenic CD45.1 mice with the BM from either CD45.2 WT or *Ifnar1^-/-^* mice (Fig 3V). After 12 weeks, 90- 95% of the recipient mice were reconstituted with the donor BM cells (SFig. 3S). Similar to *Ifnar1^-/-^* mice, β-glucan-treated WT mice reconstituted with the BM of *Ifnar1^-/-^* mice revealed no enhancement in morbidity or mortality (Fig 3W-X). These results further implicate the critical role of type I IFN signalling in the hematopoietic compartment for β-glucan-mediated protection against IAV.

### T cell-mediated immunity is required for the recruitment of β-glucan-trained neutrophils into lung tissue

Having shown the contribution of innate immune cells to the protection by β- glucan, we then investigated adaptive immunity. To determine the potential contributions of adaptive immunity in β-glucan-mediated protection against IAV, we treated *Rag1^-/-^* mice (lack T and B cells) with β-glucan 7 days prior to IAV infection. *Rag1^-/-^* mice were equally susceptible to IAV infection (lethal and sublethal dose) irrespective of β-glucan treatment (Fig. 4A-C). Interestingly, the expansion of LKS, MPP, and GMP, as well as the bias towards GP, was intact in the BM of *Rag1^-/-^* mice after β-glucan treatment (Fig. 4D-H). Furthermore, increased GP was correlated with the increased neutrophils in the BM of β-glucan-treated *Rag1^-/-^* mice (Fig. 4I and SFig. 4A). However, unlike WT mice, the recruitment of the neutrophils into the lung, spleen, and peritoneum was impaired in β-glucan-treated *Rag1^-/-^* mice (Fig.4J; SFig. 4B-F). Similarly, β-glucan-treated *Rag1^-/-^* mice failed to show an increase in the recruitment of neutrophils to the lungs post-IAV infection (Fig. 4K-M and SFig. 4G, H). This data suggests that the β-glucan promotes granulopoiesis independent of adaptive immunity, but it is required for the recruitment of neutrophils in the lung tissue.

**Figure 4.**
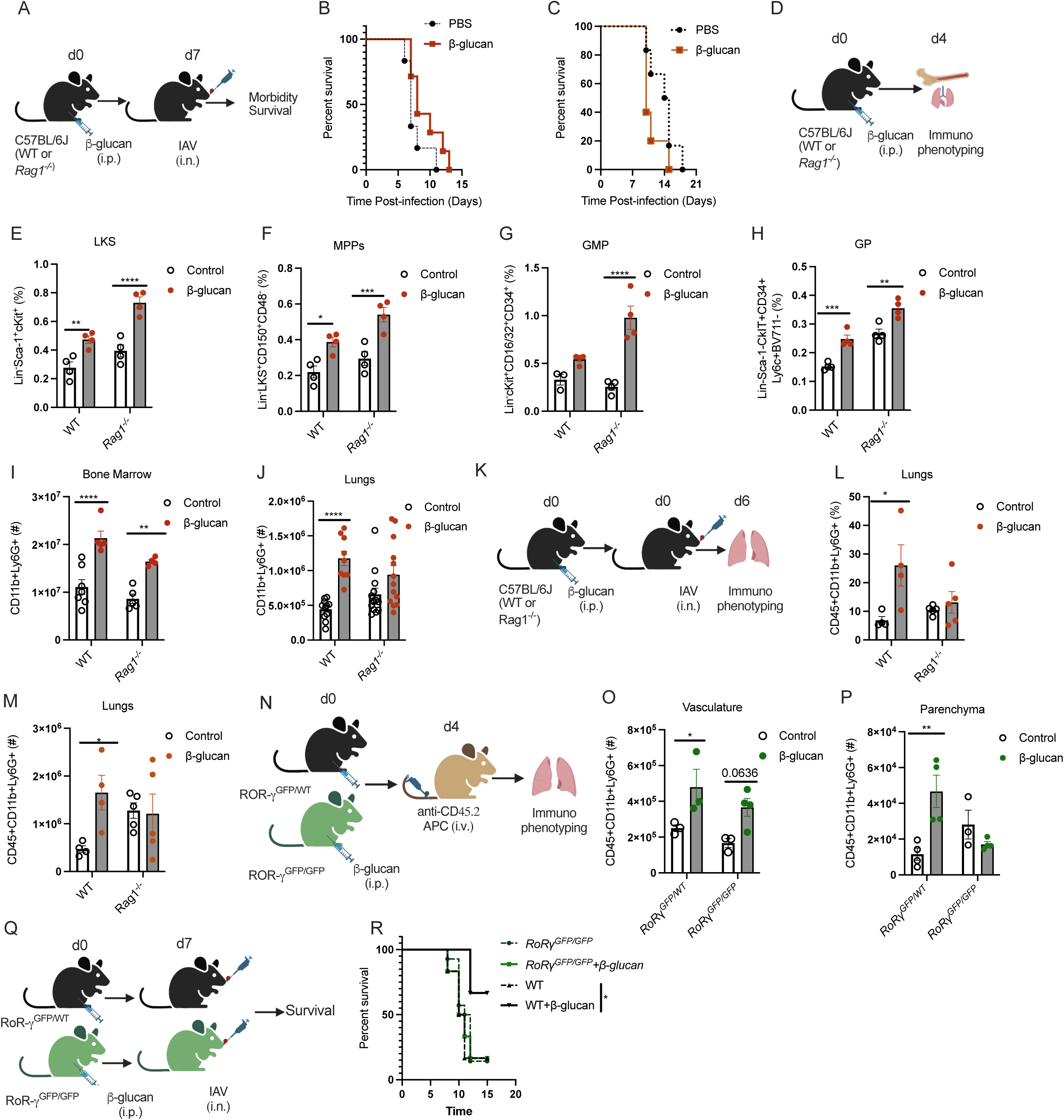
T cells are required for the recruitment of β-glucan trained neutrophils to the lung tissue. (A-C) C57BL/6 (WT and *Rag1^-/-^)* mice were infected with IAV lethal dose (B) and sublethal dose (C) at day 7 post-glucan treatment to assess survival. (D-I) C57BL/6 (WT and *Rag1^- /-^)* mice were treated with β-glucan. Expansion of LKS (E); MPP (F); GMP (G); GP (H) was assessed in the BM on day 4 post-glucan treatment. Total cell counts of neutrophils in BM (I) and lungs (J) on day 4 post-β-glucan treatment. (K-M) C57BL/6 (WT and Rag1^-/-^) mice were treated with β-glucan followed by IAV infection at day 7. Frequency (L) and total cell counts (M) of neutrophils in the lungs of β-glucan treated mice at day 6 post-IAV infection. (N-P) Intravascular staining of ROR-γ^GFP/GFP^ or ROR-γ^WT/GFP^ at day 4 post-glucan treatment. Total cell counts of neutrophils in the lungs vasculature (O) and parenchyma (P) at day 4 post-glucan treatment. (Q, R) ROR-γ^GFP/GFP^ or ROR-γ^WT/GFP^ mice were infected with IAV lethal dose at day 7 post-glucan treatment to assess survival. Data were analyzed using two-way ANOVA followed by Sidak’s multiple comparisons tests. Survival was monitored by a log-rank test. * p<0.05, ** p<0.01, *** p<0.001, **** p<0.0001.

IL-17A and IL-17F, ligands for IL-17 receptor (IL-17R), have been shown to mediate neutrophil migration into the lung in response to LPS or Gram-negative bacterial pneumonia^24, 25^. Defects in the IL-17 axis result in decreased neutrophil response associated with higher bacterial burden and poor survival of mice^26^. The RoRγ locus encodes a transcription factor required to differentiate IL-17-producing cells^27^. Thus, we treated RoRγ^GFP/GFP^ mice (lack expression of RoRγ) with β- glucan and found that similar to WT mice, the frequency and total cells of neutrophils in the BM was increased, suggesting the increase of GP in response to β-glucan was independent of IL17 (SFig. 4I). However, RoRγ^GFP/GFP^ mice failed to recruit neutrophils into the lung tissue after receiving β-glucan (Fig 4N-P). Further, β-glucan-treated RoRγ^GFP/GFP^ mice failed to show enhanced survival against IAV infection (Fig. 4Q, R). Collectively, these data suggest that β- glucan treatment promotes granulopoiesis independent of adaptive immunity and IL-17 in the BM, but RoRγ^+^ cells are required for the recruitment of neutrophils into the lung tissue.

### β-glucan reprogrammed neutrophils with regulatory phenotype and altered metabolism

Neutrophils were long considered to be terminally differentiated, homogenous effector cells. However, recent studies have demonstrated that neutrophils show a spectrum of plasticity from antimicrobial and pro-inflammatory to promote tissue healing and regulate inflammation^28^. For instance, immature and mature neutrophils have been documented not only to differ with unique surface markers but also in their functions^29^. Therefore, we next investigated whether β-glucan reprogramming of HSCs gives rise to a unique subset of neutrophils. Using multi-parameter spectral flow cytometry, we observed that β-glucan reprogramming led to a robust rearrangement of neutrophil clustering (Figure 5A-D). Under basal conditions, β-glucan shifts the whole neutrophil population toward a less activated phenotype (CD62L^high^, CD11b^low^, CD49d^low^) (Cluster 1 to Cluster 2, Figure 5B-D), with lower levels of Ly6G, CXCR2 and CD101, suggesting a lower maturation stage. Strikingly, this reshaping appears to be conserved under influenza infection, with β-glucan remaining the main driver of neutrophil clustering (Figure 5B, C). These data suggest that - even under inflammatory conditions - β-glucan reprogramming was maintained in neutrophils. Next, we investigated the impact of β-glucan reprogramming of neutrophils within IAV-infected lungs (Figure 5E-H). Under IAV infection, neutrophils infiltrate the parenchyma of the lungs and as expected, we could identify an IAV-driven population of neutrophils (Clusters 5, 6 and 8, Figure 5E, F) which represents up to a 1/3^rd^ of the pulmonary pool (Figure 5H). Despite the increased neutrophil infiltration after β-glucan treatment (Figure 2), we observed that trained neutrophils are less prone to give rise to IAV driven clusters (figure 5G, H), which are kept under 15% of the global lung populations in the infected lungs. All together, these phenotypical data show that β-glucan reprogramming give rise to a qualitative shift in neutrophil populations, which alters their behavior within the lungs under IAV infection.

**Figure 5.**
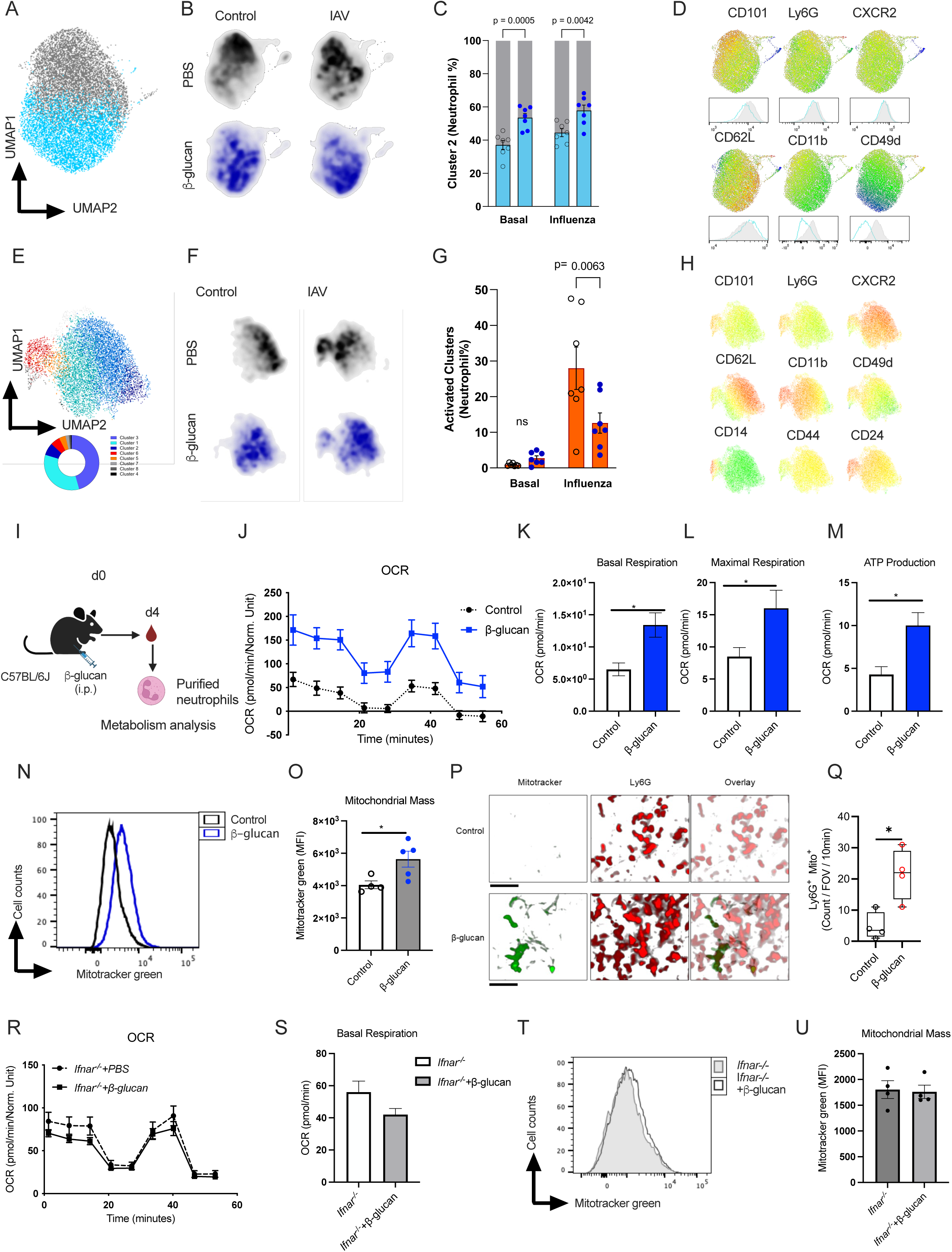
β-glucan training induces robust changes in neutrophils phenotype and metabolism. (A-D) Mice were treated with β--glucan 6 days before influenza infection. On day 9 post-glucan treatment, blood (A-D) and Lung (E-H) neutrophils from infected and non-infected mice were extracted and their surface phenotype was assessed by spectral flow cytometry. (A) Uniform Manifold Approximation and Projection (UMAP) of CD11b+ Ly6G+ neutrophils from control and β--glucan treated mice with and without influenza infection. Neutrophils separate in two main Flowsom Clusters (cluster 1, grey and cluster 2, Cyan, 96% of neutrophils) (B) UMAP from A is projected for the 4 experimental group (PBS +/- Influenza, black and β--glucan +/- Influenza, Dark Blue). Neutrophil density repartition shows a shift of neutrophils in β-glucan treated group from Cluster 1 to Cluster 2. (C) Quantification of neutrophil repartition. (D) MFIs of selected markers are projected on the UMAP from (A). Histograms show MFIs from Cluster 1 (grey and cluster 2 (Cyan). Cluster 2 neutrophils exhibit lower expression of classical maturation markers CD101, Ly6G and CXCR2 and present with a less activated phenotype (CD62Lhigh, CD11blow, CD49dlow). (E) UMAP of CD11b+ Ly6G+ neutrophils from control and β--glucan treated mice with and without influenza infection. Neutrophils separate in 8 Flowsom Clusters. (F)UMAP from A is projected for the 4 experimental group (PBS +/- Influenza, black and β-glucan +/- Influenza, Dark Blue). Clusters 5, 6, and 8 expand dramatically during influenza infection. (G) Quantification of neutrophil present in influenza-driven clusters (Clusters 5, 6, 8). (H) MFIs of selected markers are projected on the UMAP from (E). Influenza-driven clusters exhibit an activated phenotype (CD14high, CD24high CD11b high, CD62Llow, CD49dlow). MFI Mean Fluorescence Intensity. UMAPs are based on 16 surface markers (CD45, Ly6G, CD101, CXCR2, CXCR4, CD62L, CD24, CD11b, CD49d, CD44, CCR2, Ly6C, CD80, MHCII, CD16, CD14). n=7. (I-O) Mice were treated with β-glucan. Neutrophils were purified from blood on day 4 post- glucan treatment. Neutrophils’ cellular metabolism was determined by seahorse (J), basal respiration (K), maximal respiration (L), and ATP (M) production are shown. (N, O) Representative histogram plot and quantification (MFI) for mitochondrial mass using mitotracker green dye in the neutrophils of β-glucan-treated mice. (P) Representative 3-dimensional reconstructed lung intravital microscopy images from control or β-glucan treated Ly6G-TdTom mice. Mitotracker green dye was given i.v. prior to imaging. (Q) Mitochondria bright neutrophils were quantified. All scale bars = 50µm. (R-U) *Ifnar1*^-/-^ mice were treated with β-glucan. Neutrophils were purified from blood on day 4 post β-glucan treatment. Neutrophils’ cellular metabolism was determined by seahorse (R) and basal respiration (S). (T, U) Representative histogram plot (T) and quantification (MFI) for mitochondrial mass (U) using mitotracker green dye in the neutrophils from the lungs of β-glucan-treated *Ifnar1^-/-^* mice. Data were analyzed using two-way ANOVA followed by Sidak’s multiple comparisons tests and unpaired t-test. * p<0.05, ** *p*<0.01, ***p<0.001.

Cellular metabolism plays an important role in the function and plasticity of diverse immune cells, and the mechanism involved in metabolic regulations of neutrophils are continuously unfolding. Interestingly, gene expression analyses revealed several metabolic pathways (e.g., lipid and carbohydrate metabolism) that were significantly differentiated (FDR<0.01) in their response to IAV infection when comparing to non-treated with β-glucan treated mice (SFig. 5A, B), which prompted us to further investigate the potential role of metabolic changes on the functional properties of β-glucan-reprogrammed neutrophils. Under basal conditions, neutrophils are glycolytic with very few mitochondria, and thus usually, mitochondria do not contribute significantly to their metabolism ^30, 31^. However, neutrophils purified from β-glucan-treated mice rely on oxidative metabolism (Fig.5I-M). An increase in mitochondrial respiration in β-glucan- reprogrammed neutrophils was correlated with an increase in mitochondrial mass (Fig. 5N, O). To directly investigated if β-glucan induced neutrophils with increased mitochondria, Ly6G-TdTom mice were treated with either saline or β-glucan and lung intravital microscopy was performed. Mito tracker was administered i.v. to detect neutrophils with increased mitochondria. Similar to in vitro results (Fig. 5N, O), β-glucan treated mice demonstrated significantly increased numbers of Mito tracker bright neutrophils (Fig. 5P, Q). The increase of mitochondrial mass and OCR in β- glucan-treated neutrophils was also dependent on type I Interferon singling (Fig. 5 R-U), highlighting the critical role of type I IFN in reprogramming HSCs in the bone marrow (Fig. 3I- P). These results showed that β-glucan generates a subset of regulatory neutrophils with unique metabolic reprogramming, which differed from mature neutrophils.

### Survival against IAV require β-glucan-mediated regulatory neutrophils

To directly assess the role of neutrophils in survival against IAV, we depleted neutrophils (anti-Ly6G) in control and β-glucan-treated mice starting one day prior to the IAV infection and continued up to 5 consecutive days (Fig 6A). Depletion of neutrophils was confirmed in the blood of mice (S Fig. 6A). The depletion of neutrophils completely abrogated the survival advantage of β-glucan-treated mice after IAV-infection demonstrating β-glucan-trained neutrophils were essential for the host survival after the lethal infection with IAV (Fig. 6B, C). To further investigate that this enhanced survival against IAV was mediated by β-glucan-trained neutrophils and not monocytes, we performed a series of experiments using CCR2-/- mice. Similar to WT mice, treatment of CCR2-/- mice with β-glucan led to an increase in neutrophils in the BM, blood, and lungs on day four post-β-glucan treatment (S Fig. 6B-F). Importantly, CCR2-/- mice receiving β-glucan showed reduced weight loss and enhanced survival against IAV infection (Fig. 6D-F). Collectively, these data demonstrated that β-glucan increased host survival against IAV infection via reprogramming of neutrophils in the BM, which was independent of monocytes.

**Figure 6.**
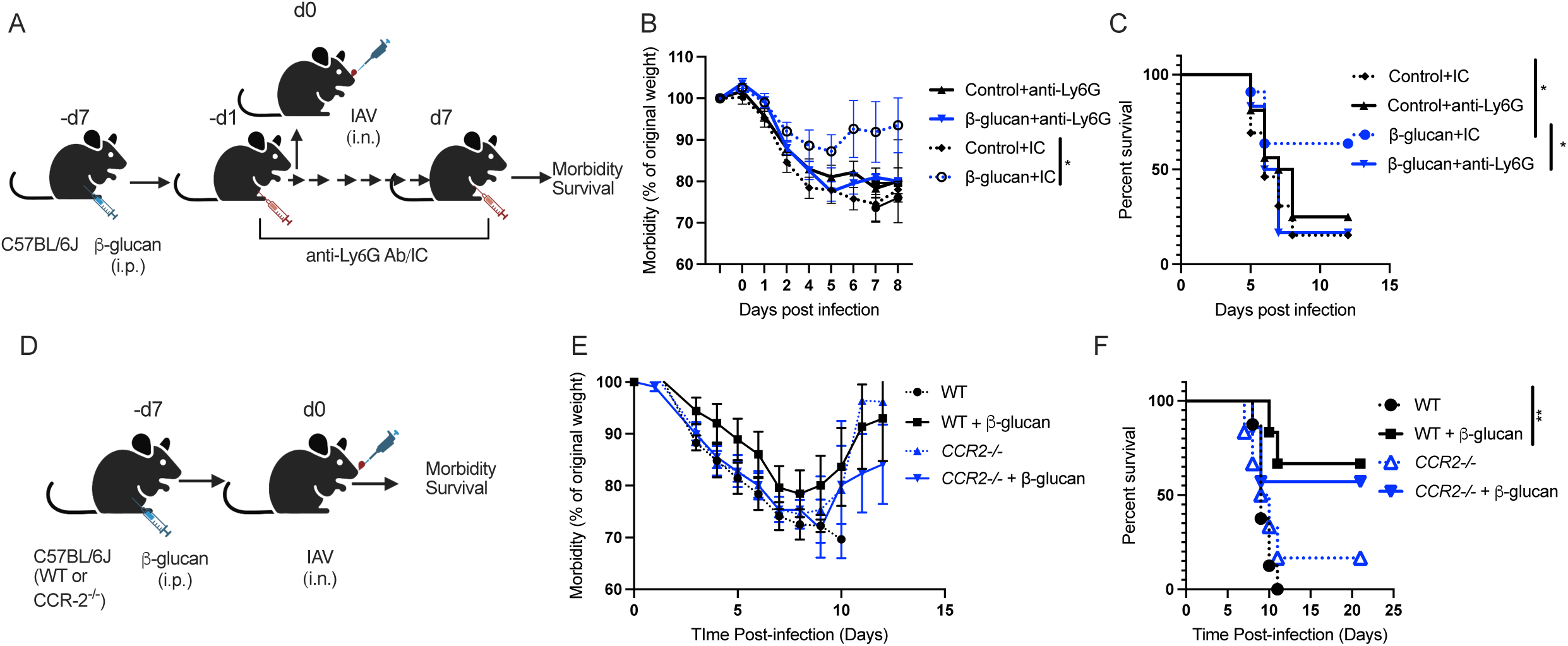
β-glucan-driven protection against IAV depends on the trained neutrophils. (A-C) Schematic of neutrophils depletion experiment using anti-Ly6G Abs. Mice were infected with IAV at day 7 post-glucan treatment. Anti-Ly6G Ab or isotype control (IC) was administered (i.p) on day-1 prior to infection and consequently until day 7. Morbidity (B) and survival (C) was monitored over time. (D-F) C57BL/6 (WT and *CCR-2*^-/-^) mice were treated with β-glucan, followed by IAV infection at day 7 post-glucan treatment. Morbidity (E) and survival (F) was monitored over time. Data were analyzed using two-way ANOVA followed by Sidak’s multiple comparisons tests. Survival was monitored by a log-rank test. * *p*<0.05, ** *p*<0.01.

## DISCUSSION

Disease tolerance is an evolutionarily conserved host defence strategy that preserves the tissue integrity and physiology without exerting a direct negative effect on the pathogen load^32^. Although the cause of severe pneumonia in the vast majority of ICU patients is due to impaired disease tolerance^33, 34^, the cellular and molecular mechanisms underlying disease tolerance remain poorly understood. In fact, our lack of understanding of immunity against infectious diseases might be due, in part, to our bias toward studying host resistance to infections and the relative lack of attention to disease tolerance.

Aberrant pulmonary innate immune responses correlate with the pathogenesis of multiple human respiratory viral infections, including IAV infection^35^. While the production of pro-inflammatory cytokines/chemokines via innate immune cells (e.g. macrophages) and the recruitment of innate immune cells are required for host resistance against pulmonary viral infections, dysregulation in cytokines production is associated with “cytokine storm”, exacerbated innate-mediated lung injury and poor clinical outcomes^36^. Thus, a controlled innate immune to pulmonary viral infections is essential for preventing excessive lung tissue pathology and preserving lung function. Here we have demonstrated that β-glucan enhances host survival against lethal infection with IAV via reprogramming HSCs towards granulopoiesis, generating a unique subset of regulatory neutrophils, which are required to promote disease tolerance against IAV infection.

The immunomodulatory activity of β-glucan depends on its physical properties (for example, particle size, molecular weight, conformation, branching frequency, or solubility)^37^. However, we also found that the strength of β-glucan signalling has a profound impact on the outcome of innate memory responses. Here we showed that one dose of β-glucan administration promotes granulopoiesis and neutrophil differentiation, with no significant difference in myelopoiesis or monocytopoiesis that we have previously observed with two doses of β-glucan against *Mtb* ^9^. Importantly, the signalling pathway involved in β-glucan-mediated granulopoiesis was dependent on type I IFN, while β-glucan-mediated myelopoiesis required IL-1β signalling^9, 11^. After one dose of β-glucan administration, Mitroulis and colleagues observed increased MPPs in the BM dependent on IL-1β signalling^20^. This is consistent with our observation that β-glucan treatment increased the LKS, MPP and GMP population. However, the production of GPs, which are downstream of GMPs was dependent on type1 IFN signalling. *Ifnar1^-/-^* mice showed expansion of LKS, MPPs and GMPs in response to β-glucan treatment but with no increase in GPs-population downstream of GMPs. Similarly, a recently published study shows that granulopoiesis in the BM after one dose of β-glucan treatment in the setting of cancer depends on the type I IFN pathway ^11^. Moreover, we speculate that the HSCs response is context-dependent and requires an additional signal. For instance, in the context of IAV infection, type I IFN was a major driver as the generation of trained GMPs/neutrophils as well as their beneficial functions were intact. This is in sharp contrast to *Mtb* infection where IL-1 signaling is dominant. The presence of pathogens^38^, pathogen- products^39^ and host mediators (e.g. cytokines)^40^ provides a complex matrix and how HSCs perceive and/or respond to these signals requires further investigations.

Although the reprogramming of HSCs/GMPs/GPs/neutrophils was entirely independent of adaptive immune cells, the recruitment of trained neutrophils into the lung required T cells. Our finding is in line with the evidence that T-lymphocytes are involved in orchestrating the sustained mobilization of neutrophils^41^. While we did not investigate the cytokine(s) involved in neutrophil recruitment into the lung, we identified that RoRγ^+^ cells were required for this recruitment. Th17 cells, γδ T cells, and type 3 innate lymphoid cells express RoRγ^+^ and can produce IL-17, which is essential for neutrophil recruitment into the lungs and protection against pathogens^42^. Moreover, the susceptibility of the β-glucan treated T and B cell-deficient host to IAV infection is an indication of the important crosstalk between trained and adaptive immunity and how both arms of immunity are required for coordinating an optimal host defense against pulmonary infections.

Conventionally, neutrophils have been considered to be a homogenous short-lived cell population, which are highly inflammatory and enhance host resistance against mainly bacterial and fungal infections^43^. However, recent studies indicate that neutrophils are a heterogenous population with a diverse array of functions from inflammatory to anti-inflammatory and pro-healing. During viral infections, neutrophils are recruited to the site of infection, but they are less directly involved in virus elimination. Our findings add to a growing body of literature demonstrating the regulatory function of neutrophils in controlling inflammation and promoting tissue regeneration and healing after tissue injury. Treatment with β-glucan reprogrammed the BM-HSCs to generate a unique subset of neutrophils with a less mature and more regulatory phenotype. Furthermore, a metabolic adaptation of neutrophils has been identified as the critical component in defining neutrophils’ functions. Conventionally, neutrophils were considered oligomitochondrial cells and their mitochondria were only associated with the cell death programs^44^. However, unlike mature neutrophils, which preferentially utilize glycolysis for energetics, immature neutrophils rely on mitochondrial respiration for energy production. It has been shown that the tumour microenvironment induces metabolic adaptation in neutrophils with increased mitochondrial fitness and immune-suppression^45^. We also found that β-glucan reprogrammed HSCs to generate regulatory neutrophils in which they maintain and utilize mitochondrial respiration for their energy demands. An additional unique feature of these regulatory neutrophils was the increased production of ROS, which has been shown to be required for polarizing macrophages towards a repair/healing phenotype during injury^46^. While we show that recruitment of β-glucan reprogrammed neutrophils is essential for host survival via promoting disease tolerance, the exact cellular and molecular mechanism(s) of how these regulatory neutrophils control inflammation or/and promote lung tissue repair requires further investigation.

The memory capacity of innate immune cells in different contexts, from sterilized inflammation to infections and cancer, is incompletely understood. While most studies on innate memory responses have been focused on the pro-inflammatory behaviour of innate immune cells and their contributions to host resistance, little is known about their potential in regulating inflammation and promoting disease tolerance. Considering the limited therapeutic options available for severe pulmonary viral infections, promoting disease tolerance via innate memory responses may be a feasible approach.

## Material and Methods

### Mice

Six- to ten-week-old C57BL/6, CD45.1, I*fnar1*-/-, IL1R-/-, Rag-/-, RoRγ-^GFP/GFP^, B6.Cg-Tg(K18- ACE2)2Prlmn/J transgenic mice were purchased from Jackson Laboratories. All animal studies were conducted in accordance with the guidelines of, and approved by, the Animal Research Ethics Board of McGill University (project ID: 5860). Mice were housed under SPF conditions with ad libitum access to food and water. Experiments were conducted using male and female sex- and age-matched mice that were randomly assigned to experimental groups.

#### β-glucan treatment

Mice were treated with β-glucan (Sigma, catalogue #G5011), 1mg/mouse i.p.

#### Viruses and Infection

All in vivo infections were performed using mouse-adapted influenza A/Puerto Rico/8/34 (H1N1) virus (IAV), kindly provided by Dr. Jonathan A. McCullers (St. Jude Children Research Hospital). Mice were challenged intranasally (in 25μL PBS) with IAV at a sublethal dose of 50 PFU or a lethal dose (LD50) of 90 PFU for the survival experiments. During survival experiments, mice were monitored twice daily for signs of duress and weighed daily. Mice reaching 75% of original body weight were considered moribund and sacrificed. Viruses were propagated and isolated from Madin-Darby Canine Kidney (MDCK) cells and titrated using standard MDCK plaque assays. MDCK cells were obtained from the American Type Culture Collection and maintained in Dulbecco’s modified Eagle medium enriched with 10% (v/v) FBS, 2 mM L-glutamine and 100 U ml−1 penicillin/streptomycin.

#### Histopathological analysis

Lungs were inflated with and fixed for 48 h in 10% formalin, then embedded in paraffin. Next, 5 μm sections were cut and stained with Haematoxylin and Eosin or Masson’s Trichrome. Slides were scanned at a resolution of 20× magnification and pictures were taken using a Leica Aperio slide scanner (Leica). Quantification of collagen-afflicted areas on Masson’s Trichrome stained slides was performed using ImageJ software (National Institutes of Health).

#### Flow Cytometry

BM/Spleen cells (3 x 106 cells) after RBC lysis were stained with fixable viability dye eFluor501 (eBioscience) at the concentration of 1:1000 for 30 minutes (4°C). Subsequently, the cells were washed with PBS supplemented with 0.5% BSA (Wisent) and incubated with anti-CD16/32 (clone 93, eBioscience) at a concentration of 1:100 in PBS/0.5% BSA at 4°C for 10 minutes except for myeloid progenitor and downstream progenitors staining. The following antibodies were then used for staining: anti-Ter-119 (clone Ter119), anti-CD11b (clone M1/70), anti-CD5 (clone 53-7.3), anti-CD4 (clone RM4-5), anti-CD8a (clone 53-6.7), anti-CD45R (clone RA3-6B2), and anti- Ly6G/C (clone RB6-8C5) all were biotin-conjugated (all BD Bioscience) and added at a concentration of 1:100 for 30 minutes at 4°C. Cells were subsequently washed with PBS/0.5% BSA. For staining of LKS, HSCs, and MPPs: Streptavidin–APC-Cy7 (eBioscience), anti-c-Kit– APC (clone 2B8, eBioscience), anti-Sca-1–PE-Cy7 (clone D7, eBioscience), anti-CD150– eFluor450 (clone mShad150, eBioscience), anti-CD48–PerCP-eFluor710 (clone HM48-1, BD Bioscience), anti-Flt3–PE (clone A2F10.1, BD Bioscience), and anti-CD34–FITC (clone RAM34, eBioscience) (all 1:100) were added and incubated at 4°C for 30 minutes. For staining of myeloid and lymphoid progenitors: Streptavidin–APC-Cy7 (eBioscience), anti-c-Kit–APC (clone 2B8, eBioscience), anti-Sca-1–PE-Cy7 (clone D7, eBioscience), anti-CD34–FITC (clone RAM34, eBioscience), anti-CD16/32 PerCP-eFluor710 (clone 93, eBioscience) and anti-CD127 BV786 (clone SB/199, BD bioscience) or anti CD127-BV605 (clone A7R34, Biolegend) (all 1:100) were added and incubated at 4°C for 30 minutes. For cMoPs and downstream progenitors: BM cells were incubated with biotin antibodies against lineage markers as mentioned above, except we included anti-Ly6G (clone 1A8-Ly6G) replacing anti-Ly6C/6G at 4°C for 30 minutes. Cells were subsequently washed with PBS/0.5% BSA. Following antibodies were added: Streptavidin–BUV- 395(BD Bioscience), anti-c-Kit–Pacific Blue (clone 2B8, BD Bioscience), anti-Sca-1–PE-Cy7 (clone D7, eBioscience), anti-CD34–FITC (clone RAM34, eBioscience), anti-CD16/32–PerCP- efluor710 (clone 93, eBioscience), anti-CD115 BV711 (clone AFS98, BioLegend), anti-Flt3–PE (clone A2F10.1, BD Bioscience), anti-Ly6C–APC (clone HK1.4, eBioscience) and anti-Ly6G AF700 (clone 1A8–Ly6G, eBioscience) all (1:100 except Streptavidin BUV395 at 1:50) were added and incubated at 4°C for 30 minutes. In some experiments cells were further stained with AnnexinV–PE and 7AAD (Biolegend) or NucSpot Far-Red (Biotium), according to the manufacturer’s instructions and unfixed cells were acquired immediately. In another set of experiments, cells were fixed and permeabilized using the FOXP3 Transcription Factor Staining Kit (eBioscience) for 1 hour at 4°C. Then, cells were stained with anti-Ki67–PE (clone 16A8, BioLegend) (1:400) for 1 hour at 4°C and acquired.

### Staining for innate and adaptive immune cells

Red blood cells were lysed in bone marrow and collagenase IV (Sigma)-treated lung samples. Lung, spleen or BM cells (3×106) were then stained with fixable viability dye eFluor501 (eBioscience) at the concentration of 1:1 000 for 30 minutes (4°C). Subsequently, the cells were washed with PBS supplemented with 0.5% BSA (Wisent) and incubated with anti-CD16/32 (clone 93, eBioscience) at a concentration of 1:100 in PBS/0.5% BSA at 4°C for 10. After washing, cells were incubated with fluorochrome tagged antibodies at 4°C for 30 minutes. Antibodies for the innate panel: anti-CD11b–Pacific Blue (clone M1/70, eBioscience), anti-CD11c–PE-Cy7 (clone HL3, BD Bioscience), Siglec-F–PE-CF594 (clone E50– 2440, BD Bioscience), F4/80–APC (clone BM8, eBioscience), Ly6C–FITC (clone AL-21, BD Bioscience), Ly6G–PerCP-eFluor710 (clone 1A8, eBioscience). Antibodies for the adaptive panel: anti-CD3–PE (clone 145-2C11, eBioscience), anti-CD19–PE-Cy7 (clone eBio1D3 (1D3), eBioscience), anti-CD4–eFluor450 or anti-CD4-FITC (clone GK1.5, eBioscience), anti-CD8– AF700 (clone 53-6.7, BD Bioscience). All cells were subsequently washed with PBS/0.5% BSA and resuspended in 1% paraformaldehyde.

### Blood Leukocytes

50 uL of whole blood collected in heparin tubes (BD) was incubated with fluorochrome-tagged antibodies at 4°C for 30 minutes. Antibodies for the innate panel: anti- CD11b–Pacific Blue (clone M1/70, eBioscience), anti-CD11c–PE-Cy7 (clone HL3, BD Bioscience), anti-Siglec-F–PE-CF594 (clone E50–2440, BD Biosciences), anti-F4/80–APC (clone BM8, eBioscience), anti-Ly6C–FITC (clone AL-21, BD Bioscience), anti-Ly6G–PerCP– eFluor710 (clone 1A8, eBioscience). Antibodies for the adaptive panel: anti-CD3–PE (clone145- 2C11, eBioscience), anti-CD19–PE-Cy7 (clone eBio1D3 (1D3), eBioscience), anti-CD4–FITC (clone GK1.5, eBioscience), anti-CD8 AF700 (clone 53-6.7, BD Bioscience). After RBC lysis, cells were subsequently washed with PBS/0.5% BSA and resuspended in 1% paraformaldehyde. If required, panels were modified to contain anti-CD45.1–APC (clone A20, BD Bioscience, 1:100) and anti-CD45.2–BUV395 (clone 104, BD Bioscience, 1:100).

Cells were acquired on the Fortessa-X20 (BD) and analyzed using FlowJo software (version 10.6.1). All percentages are of single viable frequency unless otherwise indicated.

#### Neutrophil spectral flow cytometry phenotyping

Neutrophils were harvested from mice treated with β-glucan (day0) before influenza infection (day 6), as well as from their respective controls. At day of sacrifice (day9), mice were euthanized, and blood has been harvested and kept on ice. Mice have been perfused with ice-cold PBS, lungs have been harvested, finely chopped with scissors on ice, and filtered to extract leukocytes. Blood and lungs samples were then lysed (RBC Lysis Buffer 420301, Biolegend) and kept on ice. Samples were then stained on ice with CD45-PerCP (clone 30-F11, Biolegend), Ly6G-BUV395 (Clone 1A8, BD Bioscience), CD101-APC (clone Moushi101, eBioscience), CXCR2-PE (clone SA044G4, Biolegend), CXCR4-BV711 (clone L276F12, Biolegend), CD62L-BV480 (clone MEL-14, BD Bioscience), CD24-AF700 (clone M1/69, Biolegend), CD117-BV421 (clone 2B8, Biolegend), CD11b-BUV496 (clone M1/70, BD Bioscience), CD49d-BUV563 (clone 9C10, BD Bioscience), CD44-BV570 (clone IM7, Biolegend), CCR2-BV750 (clone475301, BD Bioscience), Ly6C-BV785 (clone HK1.4 Biolegend), CD80-BV650 (clone M18/2, BD Bioscience), MHCII-BUV661(clone M5/114, BD Bioscience), CD16-PE/Dazzle594 (clone S17014E, Biolegend) CD115-AF488 (clone AFS98, Biolegend), CD14 APC/Fire780 (clone Sa14- 2, Biolegend) in staining buffer (420201, Biolegend). Samples were acquired using an Aurora spectral flow cytometer (5 lasers configuration, Cytek). Data have been analyzed using FlowJo. In brief, Cd11b^+^ Ly6G^+^ neutrophils from all mice have been concatenated to perform Uniform Manifold Approximation and Projection (UMAP) followed by Flowsom clustering using 17 surface markers (CD45, Ly6G, CD101, CXCR2, CXCR4, CD62L, CD117, CD24, CD11b, CD49d, CD44, CCR2, Ly6C, CD80, MHCII, CD16, CD14).

#### Evaluation of mitochondrial mass using Mitotracker Green

Single cell suspensions were stained with extracellular antibodies as described above and then with Mitotracker Green 150nM, (Invitrogen technologies) in PBS for 30 min at room temperature and then washed with PBS.

#### Intravascular Staining

Mice were given 2µg of FITC-conjugated anti-CD45.2 intravenously. Two minutes later, mice were euthanized, and lungs were collected and stained *ex vivo* with BUV737-conjugated anti- CD45.2 antibody in order to determine the parenchymal or vascular localization of the cells.

#### Purification of Neutrophils

Neutrophils were purified from the blood Using the EasySep^TM^ Mouse Neutrophils enrichment kit following the manufacturer’s instructions (Stem Cell Technology).

#### Extracellular Flux Analysis

Real-time oxygen consumption rates (OCR) of purified neutrophils from blood were measured in XF media (non-buffered DMEM containing 2mM L-glutamine, 25mM glucose and 1mM sodium pyruvate) using a Seahorse Xfe 96 Analyzer (Agilent Technologies). For the mitochondrial stress test, mitochondrial inhibitors oligomycin, fluorocarbonyl cyanide phenylhydrazone (FCCP), antimycin A and rotenone were used, as per the manufacturer’s recommendations. Neutrophils were purified as described above. Neutrophil adherence was achieved by plating a suspension of sorted neutrophils in seahorse assay media with 2 mM glutamine and 25 mM Glucose and spinning at the lowest acceleration to 45×*g* followed by natural deceleration. Sorted neutrophils were seeded at 0.2 × 10^6^ cells per well and incubated for 1 h at 37 °C with no CO2. XF analysis was performed at 37 °C with no CO2 using the XF-96e analyser (Seahorse Bioscience) as per manufacturer’s instructions. All measurements were normalized to cell number using a crystal violet dye extraction assay. Oxygen consumption curves were generated using Wave Desktop 2.3 (Agilent Technologies). Basal OCR was calculated by subtracting measurement 7 (non-mitochondrial respiration) from measurement 1. Maximal respiration was calculated by subtracting measurement 7 (non-mitochondrial respiration) from measurement 5, and spare respiratory capacity was the difference between maximal respiration and basal rate.

#### Intravital Microscopy

Lung intravital microscopy was performed as previously described^47^. Fluorescently conjugated anti-CD45 monoclonal antibody (5µg, clone 30-F11) was administered intravenously to discriminate intravascular leukocytes from non-intravascular leukocytes. Neutrophils were quantified using Imaris as either intravascular (TdTom+ / CD45+) or parenchymal (TdTom+ / CD45-). To visualize mitochondria high neutrophils, mitotracker green was administered intravenously just prior to imaging.

#### Ethics Statement

All experiments involving animals were approved by McGill (Permit # 2010-5860) in strict accordance with the guidelines set out by the Canadian Council on Animal Care.

#### Statistical analysis

Data are presented as mean ± SEM. Statistical analyses were performed using GraphPad Prism v9 software (GraphPad). Statistical differences were determined using two-sided log-rank test (survival studies), one-way ANOVA followed by Sidak’s multiple comparisons test, two-way ANOVA followed by Sidak’s or Dunnett’s multiple comparisons test, Student’s t-test or two-tailed Mann-Whitney test.

## SUPPLEMENTARY

### Methods

#### Bulk RNA-seq

Experimental mice were split into two groups (either infected with IAV, β-glucan treated+IAV) with 4 animals each. 7days after treatment, animals were infected with IAV and on day 6 post-infection, neutrophils were purified from the spleen using the EasySep Neutrophil Enrichment Kit and RNA was extracted using an Rneasy kit (Qiagen) according to the manufacturer’s instructions.

#### RNA-seq data processing

Adaptor sequences and low-quality score bases were first trimmed using Trimmomatic with parameters -phred33 SE ILLUMINACLIP:TruSeq3-SE.fa:2:30:10 LEADING:3 TRAILING:3 SLIDINGWINDOW:4:15 MINLEN:36^48^. The resulting reads were aligned to the mm10 mouse reference genome using STAR^49^. Read counts are obtained using featureCounts^50^ with default parameters.

#### Differential gene expression analyses

Gene expression levels across all samples were first normalized using the calcNormFactors function implemented in the edgeR R package (version 3.34.0) which utilizes the TMM algorithm (weighted trimmed mean of M-values) to compute normalization factors. Then, the voom function implemented in the limma package (version 3.38.3) was used to log-transform the data and to calculate precision-weights. A weighted fit using the voom-calculated weights was performed with the lmFit function from limma.

##### Effect of β-glucan on baseline gene expression and effect of β-glucan priming on response to subsequent influenza infection

Normalized, log-transformed gene expression levels for each sample were fit to the linear model Expression ∼ 1 + primary + secondary: primary, where primary refers to the absence of presence of β-glucan priming, and secondary refers to the uninfected or IAV infected conditions. This model captures the independent effect of β-glucan exposure on gene expression as well as the response of primed and non-primed samples to IAV infection. The make Contrasts and contrasts.fit functions implemented in limma were used to compare the gene expression response to influenza in naïve neutrophils, with that of β-glucan exposed neutrophils.

#### GSEA

Gene set enrichment analyses (GSEA) were performed using the fgsea R package (version 1.18.0) with parameters: minSize = 15, maxSize=500, nperm=100,000. To investigate pathway enrichments among genes with altered responses to IAV infection before compared to after β- glucan priming, genes were ordered by the rank statistic: -log10(pvalue)*logFC and compared with the Reactome gene sets from the MSigDB collections.

#### GO pathway enrichments and visualization

Gene ontology enrichment analyses for genes whose response to IAV infection were modulated by β-glucan priming (p < 0.05) were performed and visualized using the ClueGO application of Cytoscape^51^.

## ACKNOWLEDGEMENT.

The authors acknowledge technical help from staff at the RI-MUHC histopathology platform. This work was supported by a Canadian Institute of Health Research (CIHR) Project Grant (168885) and Bill & Melinda Gates Foundation (INV-003360) to M.D. M.D. holds a Fonds de Recherche du Québec–Santé Award and the Strauss Chair in Respiratory Diseases and is a fellow member of the Royal Society of Canada. N.K. was supported by a Fonds de Recherche du Québec– Santé Fellowship and a CIHR fellowship. O.S. receives funds from the DFG (CRC TRR332 projects A2 and Z1). R.C. receives support from the IMF of the Medical Faculty of the University of Münster. The funders had no role in study design, data collection and analysis, the decision to publish or the preparation of the manuscript.

## AUTHOR CONTRIBUTIONS

M.D. and N.K. conceived the project and designed the experiments. NK., R.C., S.S., M.S., E.P., A.G., J.D. and B.G.Y. performed the experiments. N.K., R.C., S.S., and M.D. analyzed the data. L.B. and S.S. performed RNA Seq bioinformatics analysis. N.K. and M.D. wrote the paper. M.D. supervised the project.

---

**Supp. Fig.1.**
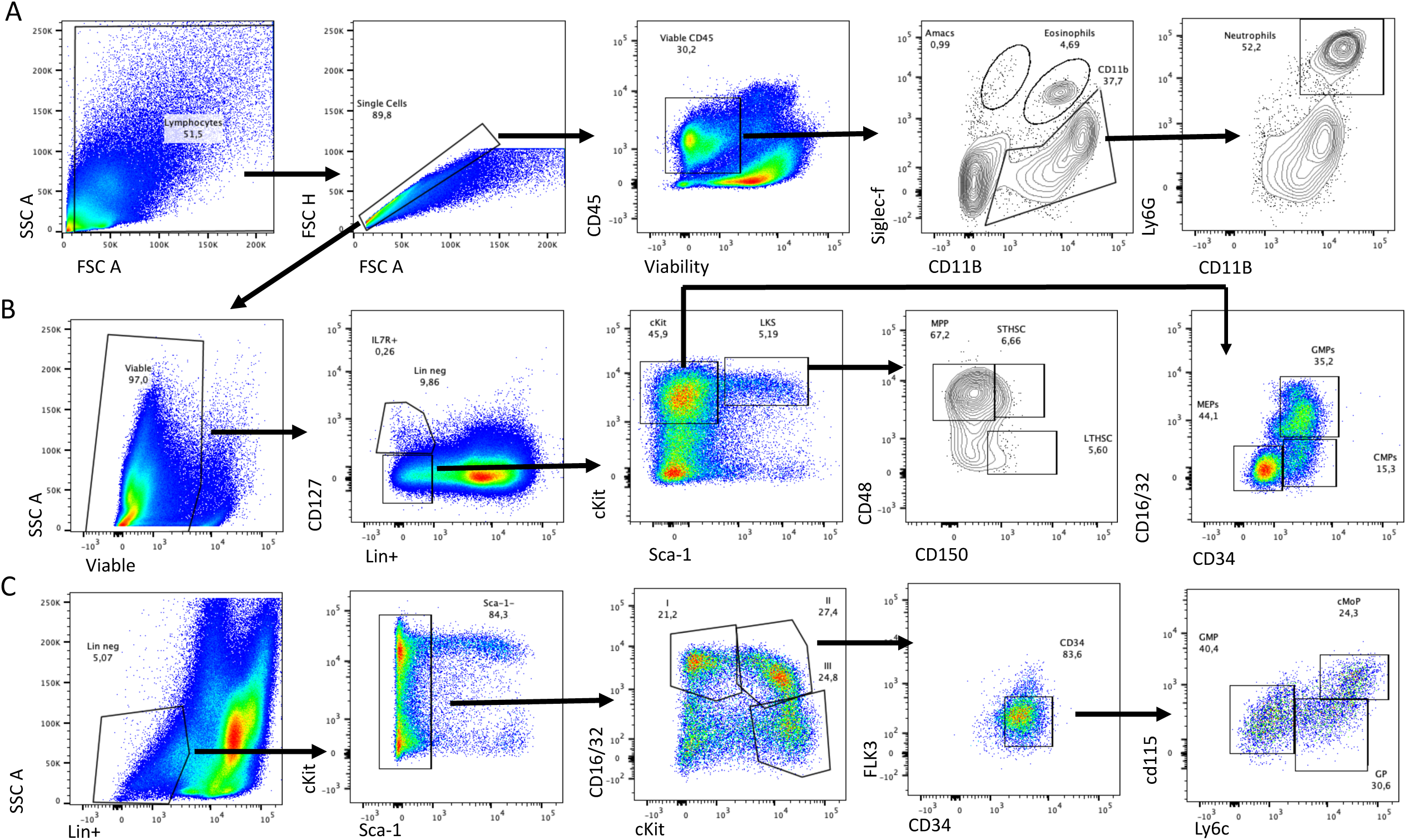
Gating strategy. (A) Cells were gated for FSC-A against SSC-A. Doublets were excluded using FSC-H against FSC-A. Viable CD45+ cells were gated, and within the viable CD45+ cells, cells were gated as CD11b, excluding SiglecF+ cells. CD11b and Ly6G double-positive cells were gated as neutrophils. (B) Cells were gated for FSC-A against SSC-A. Doublets were excluded using FSC-H against FSC-A. Viable cells were gated, and lineage-committed cells were excluded. Within the lineage-negative population, cells were gated as CD127+ and CD127-. Lin-CD127- population was further gated as LKS-defined as double positive for cKit and Sca-1, and cKit is gated as Sca- 1 negative and cKit positive. Gated on the LKS population, cells were divided into LT-HSC, ST-HSC and MPP based on CD150 and CD48 expression. C-Kit+ Sca-1- cells were further gated based on CD34 and CD16/32 to define CMP, GMP and MEP. (C) Finally, in another set of experiments, Lineage+ cells and then Sca-1+ cells were excluded. The remaining cells were subdivided into cKit+ CD16/32+ (II) and cKit+ CD16/32- groups. In the cKit+CD16/32+, cells were further gated on CD34+ Flt3- cells. Within this fraction, Ly6C+ CD115- cells were the GP, and Ly6C+ CD115+ were cMoP.

**Supp. Fig. 2.**
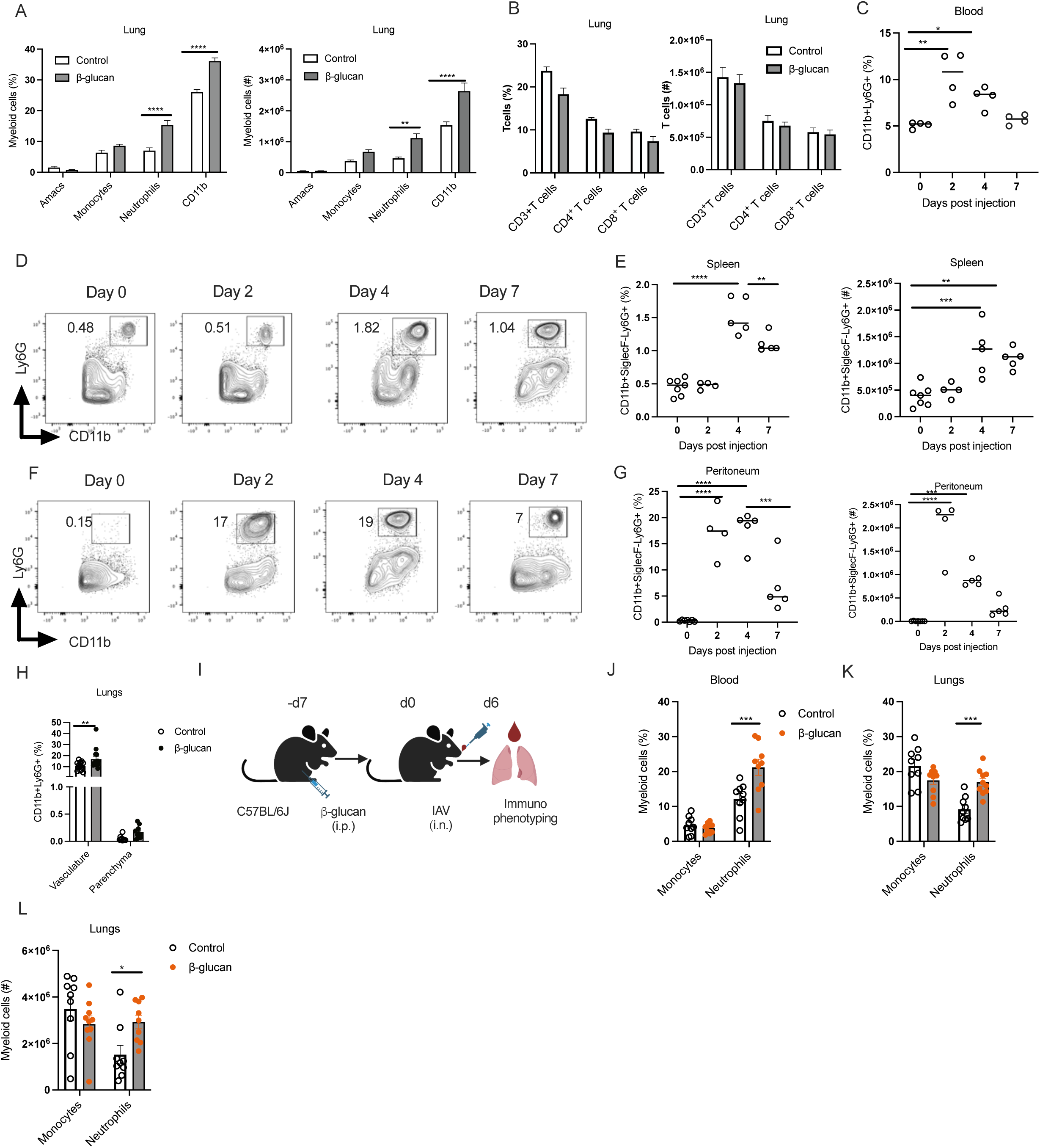
β-glucan treatment promotes granulopoiesis. **(A-P)** Mice were treated with β- glucan. Immune cells were assessed on day 4 in the lungs. Frequency and total cell count of myeloid cells (A); and adaptive cells (B) in the lungs of β-glucan-treated mice. (C-F) Kinetics of neutrophils in the blood (C) spleen (D, E) and peritoneum post-glucan treatment (F, G). The number in the FACS plot indicates the viable frequency. (H) Intravascular staining at day 4 post-glucan treatment. Frequency of neutrophils in the vasculature and parenchyma of the lungs. (I-L) Mice were treated with β-glucan, followed by IAV infection on day 7. Frequency of neutrophils in the blood (J), Frequency (K) and absolute number of monocytes and neutrophils (L) in the lungs at day 7 post-IAV infection.

**Supp. Fig. 3.**
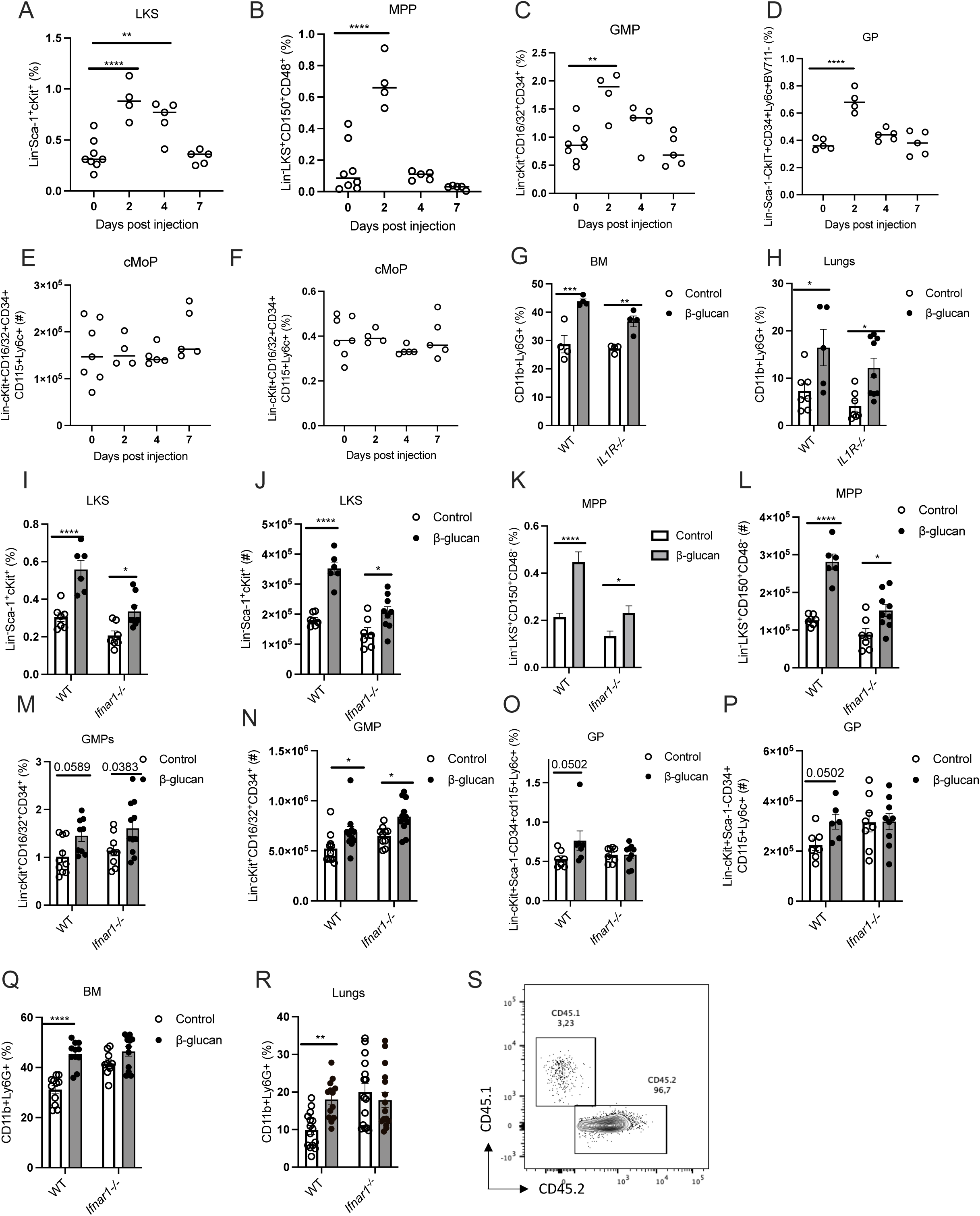
β-glucan-driven granulopoiesis is dependent on type I IFN signalling. (A-F) Kinetics of LKS/progenitors in the BM of β-glucan treated mice. Frequency of LKS (A), MPP (B), GMP (C), GP (D), cMoP (E, F) in the BM of β-glucan treated mice. (G, H) C57BL/6 (WT and IL1R^-/-^) mice were treated with β-glucan. Frequency of neutrophils in the BM (G); and lungs (H) of β-glucan treated mice at day 4. (I-R) WT and *Ifnar1^-/-^* mice were treated with β- glucan. Frequency and total cell counts of LKS (I, J); MPP (K, L); GMP (M, N); and GP (O, P) in the BM at day 4 post-glucan treatment. Frequency of neutrophils in the BM (Q) and lungs (R) at day 4 post-glucan treatment. (S) Chimerism was confirmed *via* flow cytometry.

**Supp. Fig.4.**
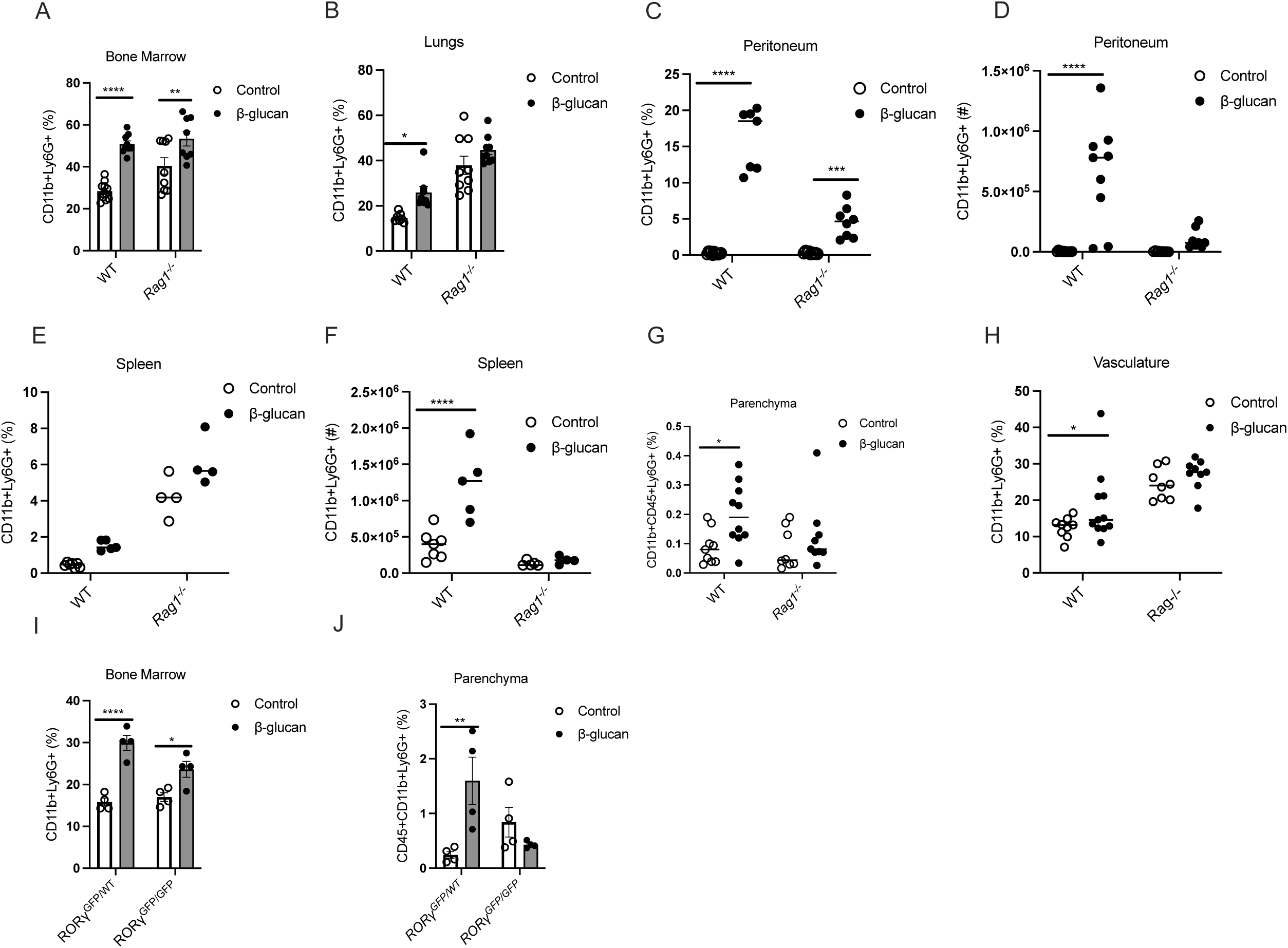
Localization of trained neutrophils is dependent on adaptive immune cells. (A-I) C57BL/6 (WT and Rag1^-/-^) mice were treated with β-glucan. Frequency of neutrophils in the BM (A) and lungs (B); frequency and absolute count of neutrophils in the peritoneum (C, D); spleen (E, F) at day 4 post-glucan treatment. Frequency of neutrophils in the parenchyma (G) and vasculature (H) of lungs at day 4 post-glucan treatment. (I, J) ROR-γ^GFP/GFP^ or ROR-γ^WT/GFP^ mice were treated with β-glucan. Frequency of neutrophils in the BM (I); and lung parenchyma (J) at day 4 post-glucan treatment.

**Supp. Fig. 5.**
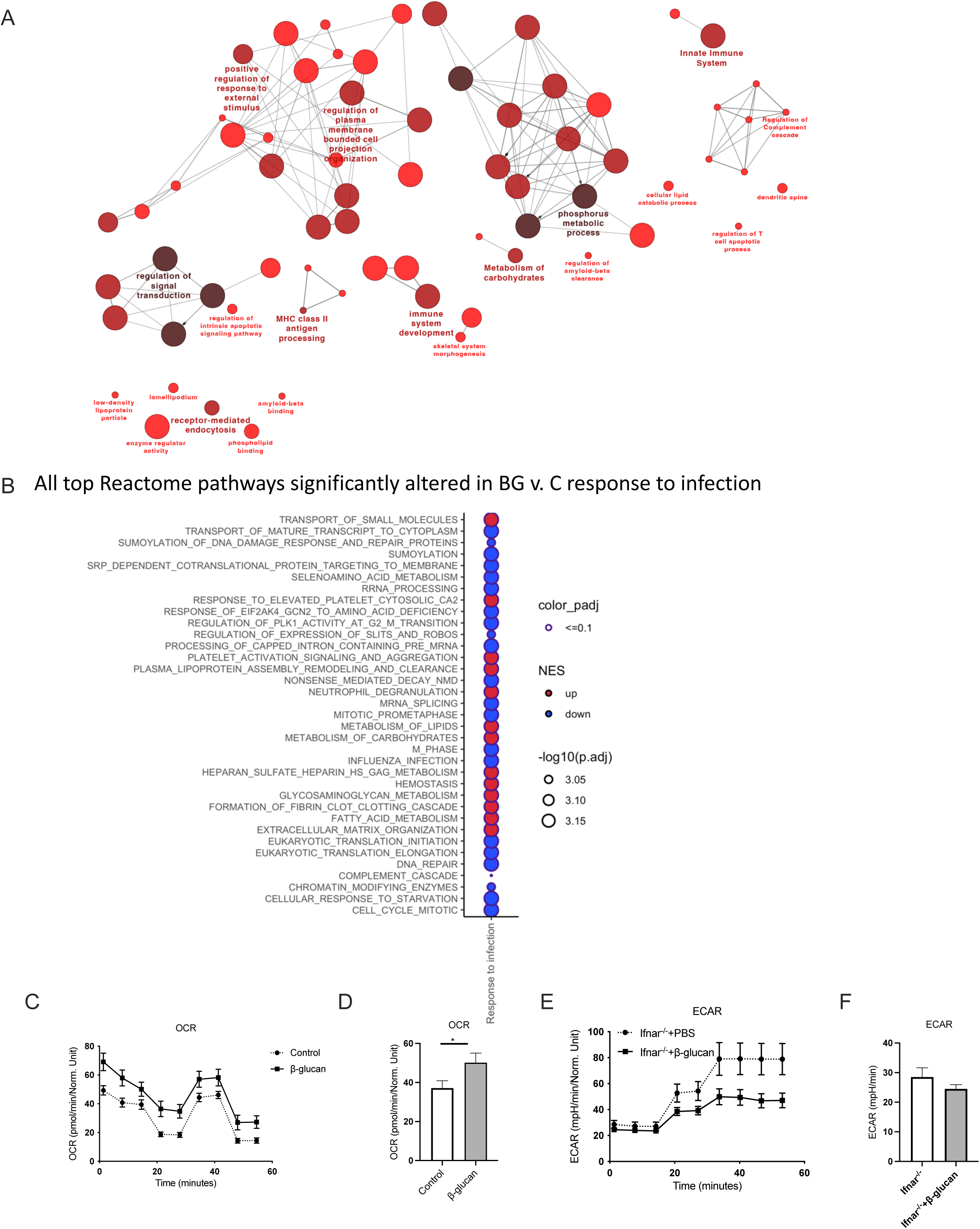
β-glucan treatment augments mitochondrial respiration. (A-F) Mice were treated with β-glucan, followed by IAV infection on day 7. Splenic neutrophils were purified at day 4 post IAV and subjected to RNA Seq (A, B), and purified blood neutrophils were subjected to seahorse metabolism (C-F). (A) Network visualization of GO-terms significantly enriched among genes whose response to IAV was primed by β-glucan exposure (p < 0.05). Each circle represents an enriched GO term (p.adj < 0.05). The larger nodes with darker shading indicate a greater degree of significance. Connections between nodes indicate similarity between terms. (B) Summary bubble plot of gene set enrichment analysis (GSEA) results. Genes were ordered by the rank statistic –log10(pval)*logFC for the effect of β-glucan priming on response to IAV infection and compared against Reactome gene sets. Circle size and shading is scaled to the normalized enrichment score (NES). All circles with a dark border have padj<=0.1. Red circles indicate that the pathway is increased in β-glucan primed samples and blue circles indicate that β- glucan -priming leads to an overall decrease in expression of the pathway.

**Supp. Fig. 6.**
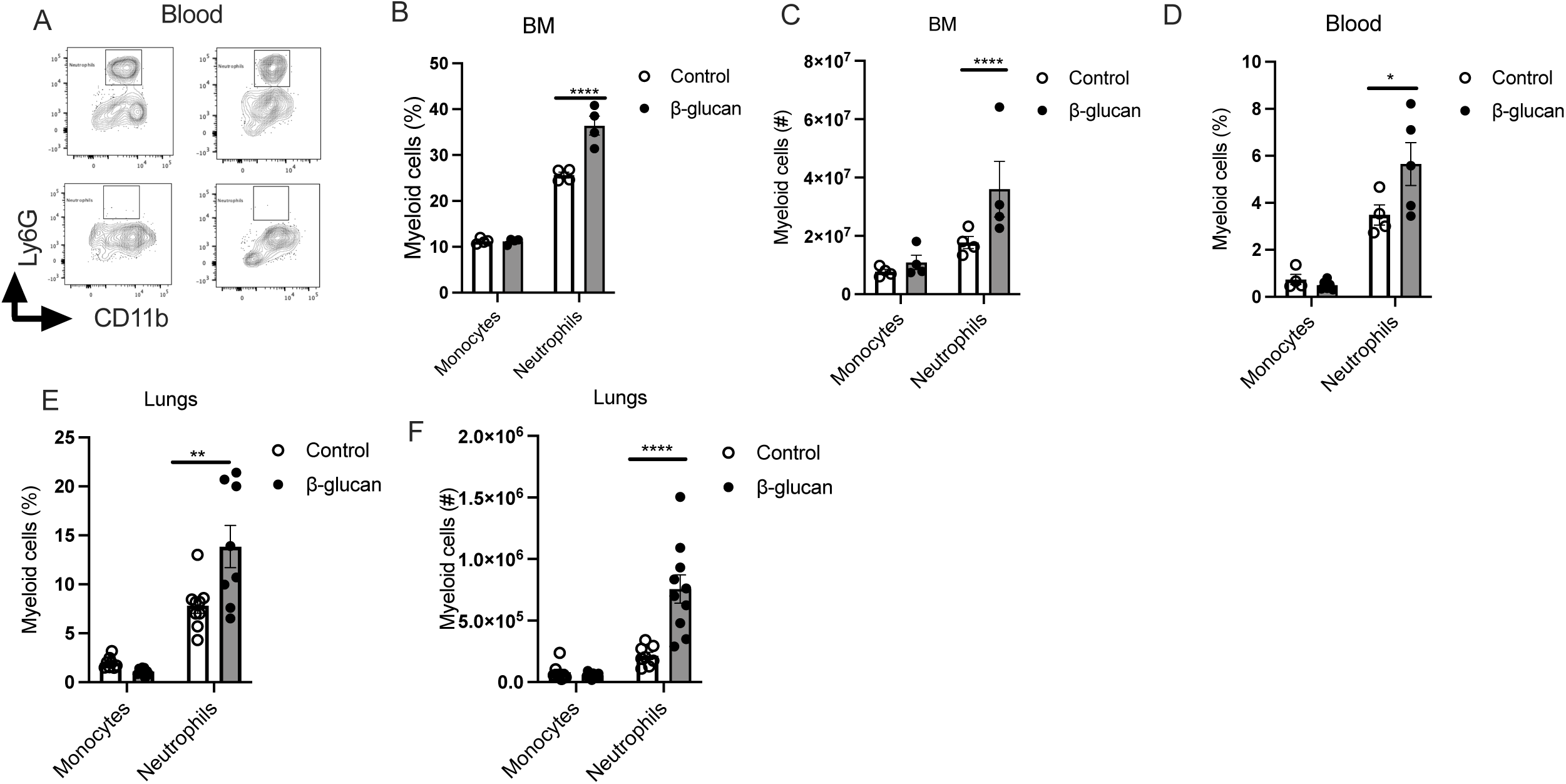
β-glucan mediated protection depends on trained neutrophils. (A) FACS plot showing the depletion of neutrophils in the blood of mice treated with anti-LY6G depletion antibody. (B-F) WT and *CCR-2*^-/-^ mice were treated with β-glucan. Frequency and absolute number of monocytes and neutrophils in the BM (B, C); blood (D); and lungs (E, F) at day 4 post-glucan treatment.

